# Protein sequence domain annotation using a language model

**DOI:** 10.1101/2024.06.04.596712

**Authors:** Arpan Sarkar, Kumaresh Krishnan, Sean R. Eddy

## Abstract

Protein domain annotation underlies large-scale functional inference and is commonly performed by scanning sequences against libraries of profile hidden Markov models (profile HMMs). We describe PSALM, a protein domain annotation method that combines (i) a pretrained protein language model (ESM-2) with (ii) a per-residue domain-state classifier and (iii) a structured probabilistic decoder that produces a single, non-overlapping set of domain calls with explicit boundaries and scores. On a benchmark of 89M protein sequences with 107M annotated domains, PSALM attains a domain-detection sensitivity-specificity tradeoff comparable to HMMER. We characterize sequence and residue-level coverage on UniProtKB, observing higher coverage for HMMER at stringent expected false positive counts (E-values) and higher coverage for PSALM at relaxed E-values. We release code for data processing, training, and inference, along with the model weights and datasets used for training, validation, and benchmarking.

## Introduction

Proteins are largely composed of distinct structural and functional units conserved through evolution, known as domains. Locating and characterizing these domains within a protein sequence, known as protein sequence annotation, is an important task in computational molecular biology — insight into the individual functions of these domains, which may act independently or in concert with neighboring domains, helps shed light on the overall biological role of the protein. As the size of protein sequence databases continues to grow rapidly, scaling to billions of sequences [1, 2, 3], so too does the number of proteins with unknown function. In order to exploit this wealth of information, we want to maximize the power of large-scale sequence annotation tools to detect domains in uncharacterized proteins and further uncover clues about the molecular basis and evolutionary trajectory of life.

The state of the art in protein domain sequence annotation uses profile hidden Markov models (profile HMMs) to detect domains [4] and profile/profile comparison to identify homologous domains [5]. Domain annotation is accomplished by scanning sequences against libraries of curated domain models. InterPro is a consortium of protein domain models from several member databases [4], and InterProScan allows sequences to be scanned against InterPro’s member database models [6]. Most member databases contributing domain models to InterPro use profile HMMs built and searched with HMMER [7], including Pfam, SMART, NCBI-fam/TIGRFAM, SUPERFAMILY, PANTHER, and CATH-Gene3D [8, 9, 10, 11, 12, 13]. Pfam, for example, currently defines ∼24,000 curated domain families. Rather than comparing a query protein sequence to hundreds of millions of sequences, annotation with respect to Pfam reduces to comparing the query against a fixed library of 24K family profile HMMs to identify domains.

Profile HMMs are created from multiple sequence alignments (MSAs) of homologous subsequences. Aligned consensus columns define a sequence of “match” states, while “insert” and “delete” states model insertions and deletions around those columns [14]. Profile HMMs have enabled sensitive, high-coverage, large-scale protein sequence annotation, but they rely on several simplifying assumptions: independence of residues given the hidden state, affine gap costs, and treating sequences as independent observations without explicitly modeling shared evolutionary history. These assumptions limit how much correlation between residues, such as conservation patterns across multiple MSA columns or domain–domain co-occurrence, can be exploited. Recent breakthroughs such as AlphaFold2 [1] and ESMFold [15] suggest that domain-annotation approaches could benefit from models that explicitly capture relationships between positions within and across domains, rather than relying only on position-wise consensus.

Most work to date on applying deep learning methods to protein sequence annotation has targeted sequence-level labels. ProtCNN and ProtENN use deep convolutional models to assign Pfam family labels to whole sequences [16]; ProtTNN frames the task with transformer-based protein language models (pLMs) [17]; and ProtEx augments a transformer with database retrieval capabilities to improve sequence-level functional annotation [18]. These methods do not return start/stop coordinates for domains. In contrast, Res-Dom [19] and DistDom [20] are segmentation models that identify domain boundaries within a sequence, but they do not predict the domain families of the resulting segments. Relying on sequence-level labels, rather than domain annotations which have explicit boundaries, risks a “transitive annotation catastrophe” [21] — if a multi-domain protein is labeled with a function because it contains domain A, that label can be transferred to homologs sharing only an unrelated domain B, propagating and amplifying misannotation. Domain-annotation deep-learning models such as ProtENN2 [22] and InterPro-N [4] have been used in practice to annotate domains and expand Pfam coverage, with predictions integrated into InterPro, and do not currently have publicly released training code or models.

Here we describe PSALM (“Protein Sequence Annotation using a Language Model”), which builds on the ESM-2 pLM with a Pfam-wide per-residue classifier and a probabilistic domain decoder that produces a non-overlapping set of domain calls with explicit boundaries for an input sequence. In domain-level benchmarks, PSALM attains sensitivity and specificity comparable to HMMER while also expanding the coverage of large-scale sequence databases, suggesting that PSALM may be a practical alternative for large-scale protein sequence annotation.

## Results

PSALM is composed of three parts (Fig. 1A): (i) An existing pLM (ESM-2) that we fine-tune, which produces per-residue contextual embeddings, or fixed-length vector representations. We hypothesize that these embeddings are sufficiently informative that domain membership is identifiable at each position individually. (ii) A domain-state classifier that converts each embedding into a per-residue probability distribution over domain states. (iii) A domain decoder that uses a structured domain-state model to turn these per-residue probabilities into well-defined, non-overlapping domain calls with boundary coordinates and confidence scores.

**Figure 1:**
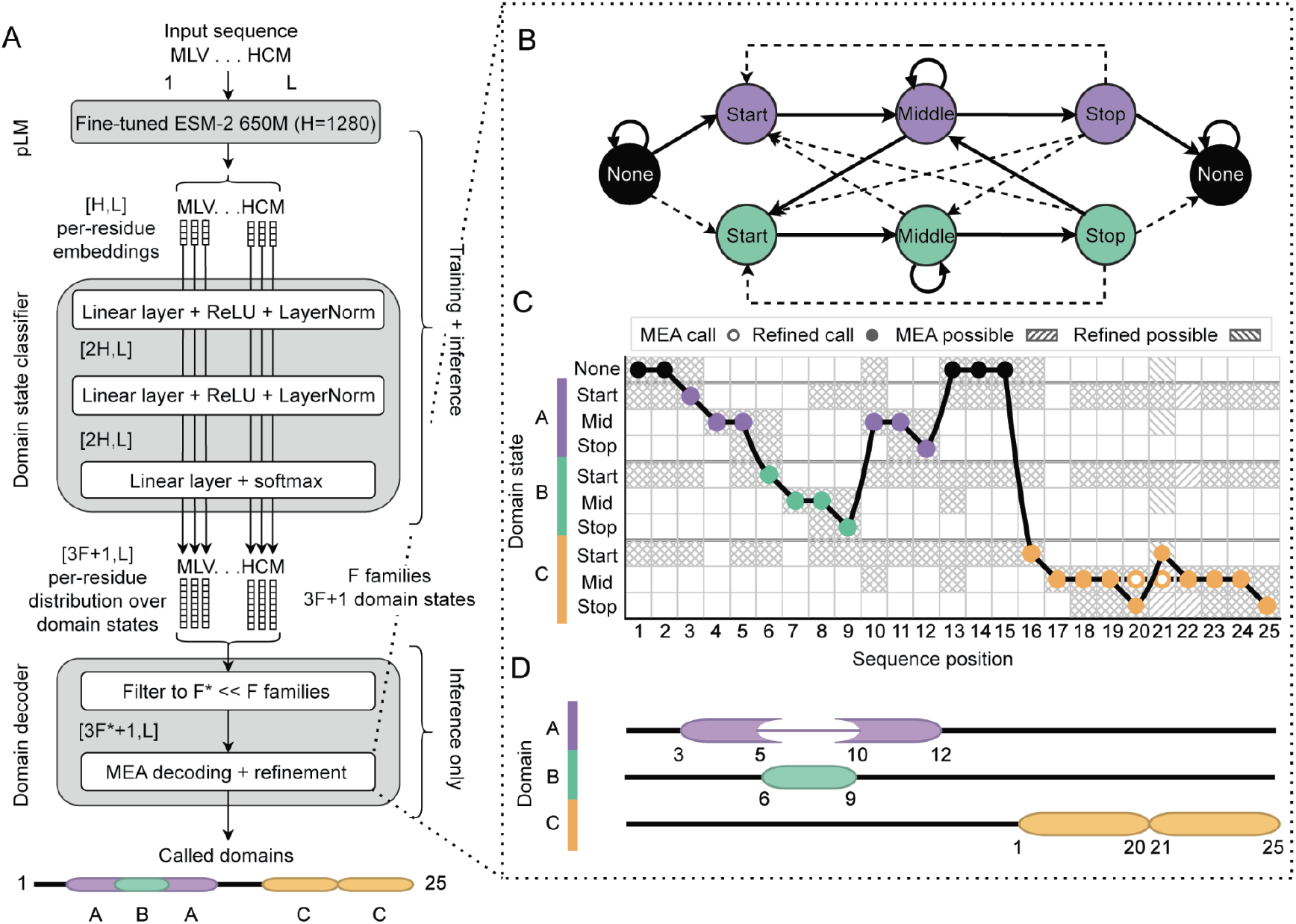
Overview of PSALM inference. The illustrated sequence (length L=25) is a toy example; real proteins containing four domains are longer. (**A**) Schematic of the three-stage pipeline: a protein language model (pLM) produces per-residue embeddings, a per-residue domain-state classifier assigns perresidue probabilities over Pfam domain states, and a structured probabilistic decoder converts these into non-overlapping domain calls with scores. (**B**) Domain state model describing the transitions within and between family A (purple), family B (green), and None. For visual clarity, the None state is duplicated but represents the same state, and the other possible cross-family transitions to family C (yellow) are excluded from this schematic. Dashed transition arrows represent paths not taken in the final call. (**C**) Two-pass maximum expected accuracy (MEA) decoding. A first pass runs forward–backward and MEA over all families, yielding an initial set of non-overlapping calls. We also show the plausible competing states at each position under the posterior. Calls flagged as over-extended or merged (e.g., two nearby family-C domains merged into one) are refined by re-decoding the region with a family-C-specific domain state model and MEA, producing the refined call and its plausible refined states. (**D**) Example domain annotation result after refinement.

**Figure 2:**
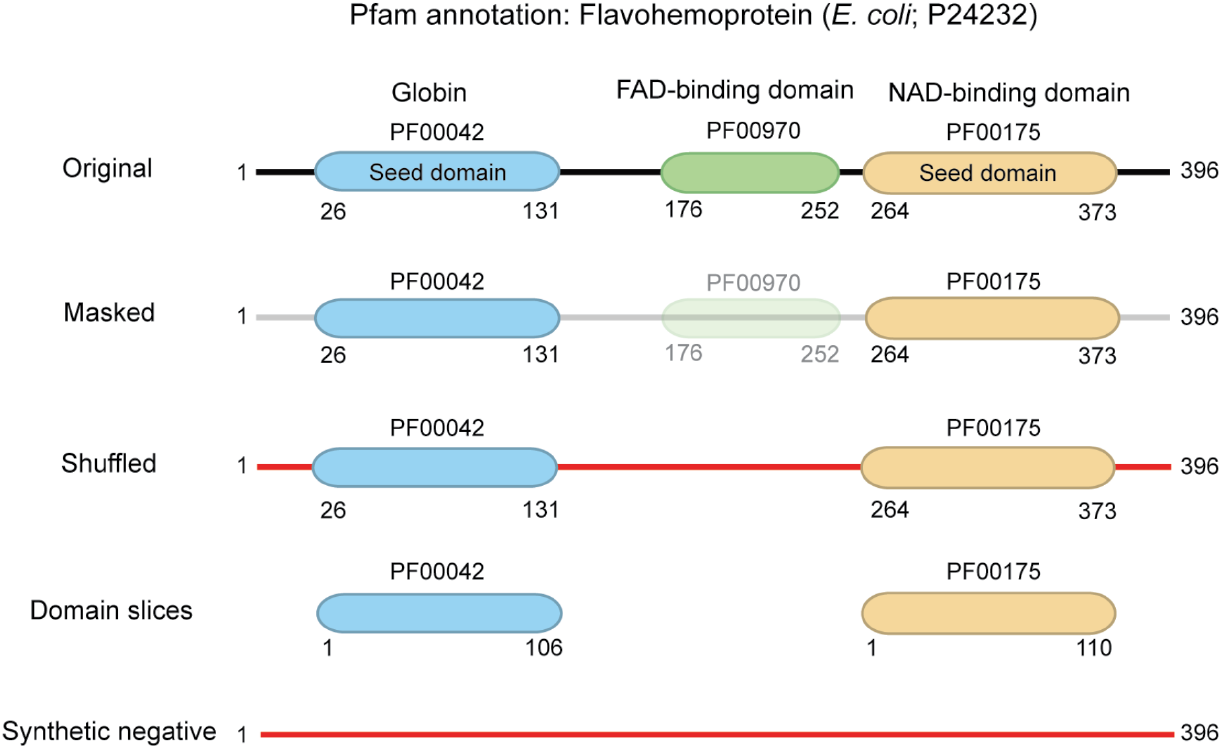
Training data augmentation. From each training set protein we generate: (i) the masked sequence (loss calculated only on unmasked domains), (ii) a shuffled-outside variant, (iii) one domain-slice per seed domain, and (iv) fully shuffled negatives labeled None. Grey indicates positions masked from the loss; red indicates shuffled residues. This example sequence has three domains, two of which are seed domains, to illustrate that a single sequence can contribute multiple domain slices; in training set 1, the average number of seed domains per sequence is 1.13, so labeling is typically sparser than in this example.

### ESM-2

We use ESM-2 650M, an openly released encoder-only protein language model that maps an amino-acid sequence *x*_1:*L*_ to contextual per-residue embeddings *h*_1:*L*_, where each *h*_*t*_ ∈ ℝ^*H*^ summarizes the sequence context around position *t* [15]. We use the 650M-parameter model as it is the largest model that remains feasible for end-to-end fine-tuning on long sequences given our resources (see Methods). Prior analyses suggest the 650M model already encodes sequence-similarity signal in its sequence-level representations that is broadly comparable to the 3B parameter model [23].

### Domain state classifier

PSALM maps an input amino-acid sequence *x*_1:*L*_ = (*x*_1_, …, *x*_*L*_) to a sequence of contextual residue embeddings using the ESM-2 650M protein language model [15] that we fine-tune. Each embedding is passed through a three-layer ***∼***200M-parameter MLP head (ReLU activation + LayerNorm in the hidden layers), producing a categorical distribution over the domain-state set 𝒮 at each position. After training, the domain-state classifier outputs a length-*L* matrix of per-residue state probabilities

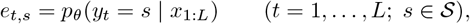

which serve as inputs to the structured decoder (Fig. 1C).

#### Data

We train the classifier with per-residue cross-entropy against full length protein sequences annotated with ground-truth domain-state labels *y*_1:*L*_ over 𝒮 (defined above). Training data are derived from two sources — one smaller, more reliably annotated database (training set 1) and one larger database (training set 2). We found that training PSALM first on training set 1 and then training set 2 results in both higher sensitivity and specificity.

We construct training set 1 from protein sequences in UniProt/UniParc release 2025 01 [4, 24]. To restrict this set to sequences containing curated domain annotations, we only retrieve the 1.2M sequences used to create Pfam-37.2 seed alignments [8] — a curated multiple sequence alignment for each of the 24K Pfam domain families. This set spans 596.3M residues of which 204.2M are labeled, totaling 1.38M seed domains - the curated subset from all detectable domains in these 1.2M sequences. During training, all unlabeled residues are treated as None (background). The validation set is sampled from UniProt sequences with Pfam domains identified by HMMER profile HMMs with curated, family-specific gathering thresholds (GAs) and does not overlap with training set 1. A random sample may be biased towards overrepresented domains in UniProt. For example, randomly sampling from a set of 10 sequences with domain A and 3 sequences with domain B will yield a set where domain A is overrepresented. To avoid such bias in the validation set, we sample 5 domains from each of the 24K Pfam domain families and retrieve their corresponding full length sequences, resulting in a validation set of 111.9K sequences with 58.2M total residues (30.1M annotated residues across 237.7K domains).

Training set 2 is also sampled from UniProt. We cluster all UniProt sequences at 30% identity to filter out closely related sequences with MMseqs2 `linclust` [25] and select the longest sequence per cluster, resulting in 28.5M sequences, totaling 11.2B residues. By restricting this set to sequences with at least one Pfam annotation, we obtain the final training set 2 of 24M sequences (9.5B residues; 36.7M domains spanning 4.6B annotated residues). This set is both larger and more densely annotated (1.51 annotated domains per sequence on average in set 2 vs. 1.13 in set 1).

#### Data augmentation

Training sequences may contain real but unannotated domains outside the labeled regions. We aim to avoid penalizing true domain predictions against partial ground truth while also exposing the model to realistic non-domain background; additionally, augmentation increases effective training diversity beyond the raw corpus size. We therefore apply an augmentation scheme to both training sources. For training set 1, each full-length sequence generates: (i) a *masked* example, where loss is computed only on unmasked domains; (ii) a *shuffled* example, where residues outside annotated domains are randomly permuted (domains unchanged) and loss is computed across the full sequence; (iii) one *domain-slice* example per annotated domain (the domain excised from its context); and (iv) *negative* examples formed by fully shuffling sequences and labeling all positions as None. This yields two augmented training set 1 corpora that differ only in the fraction of negatives: 4.0M sequences at 5% negatives and 5.0M sequences at 23.7% negatives (chosen to match the total number of sequences in the UniProtKB Reference Proteomes not annotated by Pfam [8]). For training set 2, we reduce the number of examples generated by a sequence, as training set 2 is larger and more densely annotated than training set 1: each sequence contributes either an original or shuffled-outside example (50/50), 10% of examples (3.5M) are domain slices, and negatives comprise 23.7% of the augmented corpus (8.3M, for 35.1M sequences total).

#### Training

Training proceeds by gradient-based optimization in three stages. First, we freeze ESM-2 and train only the MLP head for 5 epochs (peak learning rate 5 ×10^−4^ for head) on the training set 1 augmented corpus with 5% negatives (∼120K steps). We then unfreeze ESM-2 and continue training for 5 epochs (peak learning rates: 1× 10^−5^ for ESM-2 and 1 ×10^−4^ for head) on the training set 1 augmented corpus with 23.7% negatives (∼150K steps), using a smaller learning rate for the ESM-2 parameters than for the MLP classifier. A higher percentage of negatives in the training data helps boost specificity, especially when the ESM-2 parameters are unfrozen. To further increase the sensitivity of model using a larger dataset, we extend training using the augmented training set 2 for 1 epoch (∼220K steps, peak learning rates: 1 ×10^−5^ for ESM-2 and 1 ×10^−4^ for head). Throughout training, performance on the validation is evaluated over evenly spaced checkpoints to check for over-fitting. Full optimization hyperparameters are given in Methods.

### Structured domain-state model

For an amino-acid sequence *x*_1:*L*_, PSALM’s domain-state classifier predicts a label *y*_*t*_ ∈ 𝒮 at each position *t*, where

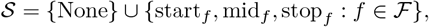

with ℱ the set of Pfam families (|ℱ| = 24,076 in Pfam 37.2) and |𝒮|= 3|ℱ| + 1 = 72,229 distinct states (Fig. 1B). The None state represents background (i.e., residues treated either as not homologous to any Pfam family or not labeled in the training corpus). For each family *f* the triple (start_*f*_, mid_*f*_, stop_*f*_) encodes the within-family structure (entry, interior, exit) of a domain instance of *f*; expected domain length is controlled by the transition probabilities out of mid_*f*_ (primarily to stop_*f*_).

We model the progression of these states along the sequence with a transition matrix

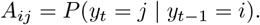

We estimate transition probabilities from labeled sequences with empirical frequencies of adjacent label pairs in the training corpus. Each row is normalized to sum to 1. Assuming domains are at least 3 residues long (start and stop with a single middle state in between), 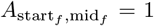 and 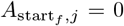 for all *j* ≠ mid_*f*_ ∀*f ∈* ℱ. Within family *f*, 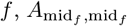 is the fraction of observed transitions out of mid_*f*_ that remain in mid_*f*_, and 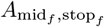 is the fraction that exit to stop_*f*_ (these two probabilities sum to 1 when no other outgoing transitions from mid_*f*_ are present). Similarly, the stop-row is estimated from counts over all observed outgoing transitions from stop_*f*_. If the training corpus does not have transitions from stop_*f*_ to an adjacent or embedded/nested domain, which we call “cross-family transitions”, then 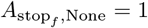. If the training corpus contains cross-family transitions, those transitions will appear as nonzero entries in *A* through the count-based estimate above.

For each input sequence of length *L*, we set the None row of *A* to impose an expected background run length of *L*:

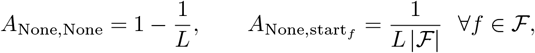

and *A*_None,*j*_ = 0 for all other states *j* (in particular 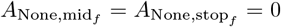), so domains can only begin in start_*f*_.

Many biologically-plausible cross-family transitions may not be observed in the training data. The domain-state model should allow for such transitions. However, the space of possible cross-family transitions is enormous and dominates the transition structure. If we were to include all allowed transitions over the full family set ℱ, then the transition matrix over domain states would contain

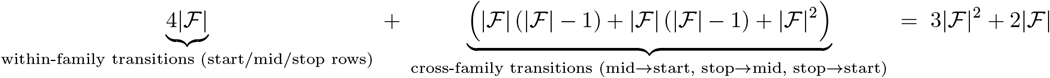

nonzero entries (for Pfam 37.2, |ℱ|= 24,076, this equals 1,739,009,480 nonzero transitions out of ∼5.2B possible transitions). Thus the cross-family terms account for *>* 99.99% of all nonzero transitions, yet only a tiny fraction are observed in our training data (e.g., *<* 0.004% of all possible cross-family transitions).

To allow unobserved but plausible cross-family transitions without considering the full |ℱ|^2^ space, we introduce three transition-mass parameters (estimated from UniProt sequences with Pfam annotations; see Methods) that allocate total probability mass to the three cross-family transition classes mid→start, stop→mid, and stop→start. For each affected source row, we distribute the corresponding mass uniformly over cross-family destinations of the relevant type that are unobserved in the training corpus, then renormalize the row to sum to 1. We apply these mass adjustments only after inference-time family filtering, when decoding is restricted to a much smaller per-sequence candidate family set.

### Domain decoding and refinement

Given the per-residue state probabilities *e*_*t,s*_, we convert noisy residue-wise predictions into a consistent set of non-overlapping domain calls (Fig. 1D) by structured decoding in a linear-chain model over the domain-state alphabet 𝒮. Using *e*_*t,s*_ as per-position state scores and the fixed transition matrix *A* defined above, we define a decoder posterior *P* (*y*_1:*L*_ |*x*_1:*L*_) over valid state paths. We compute posterior marginals by forward–backward and then identify a single path with maximum expected accuracy (MEA) decoding. This procedure is analogous to inference in a linear-chain conditional random field (CRF) [26], but here the transition structure is fixed from annotation statistics rather than learned by CRF training.

#### Inference-time family filtering

To make decoding tractable, we apply a per-sequence candidate-family filter before forward–backward/MEA. For each position *t*, we take the top-scoring non-None state *s*^*∗*^(*t*) = argmax_*s∈*𝒮\{None}_ *e*_*t,s*_ and collect its Pfam family. This yields a candidate family set ℱ^*∗*^ = {family(*s*^*∗*^(*t*)) : 1 ≤ *t* ≤ *L*} and a sub-alphabet 𝒮_sub_ = {None}∪⋃ _*f ∈*ℱ_*∗* {start_*f*_, mid_*f*_, stop_*f*_}. We then restrict both emissions and transitions to 𝒮_sub_: at each position we keep *e*_*t,s*_ only for *s ∈* 𝒮_sub_ and renormalize over the retained states, and in *A* we drop destinations outside 𝒮_sub_ and renormalize each affected row. Any probability mass removed from a row is absorbed into a single default transition (self-loop *i* →*i* for middle states and *i* →None for stop states). Finally, we overwrite the None row using the sequence-specific dwell model with starts restricted to ℱ^*∗*^, preserving an expected None run length of *L* while limiting domain entry to candidate families. On the validation set, this filter has a negligible effect on accuracy (*<* 0.1%) while substantially reducing the state space (full details in Methods).

#### Forward–backward with beam pruning

Let *α*_*t*_(*j*) = *P* (*x*_1:*t*_, *y*_*t*_ = *j*) be the forward quantity and *β*_*t*_(*i*) = *P* (*x*_*t*+1:*L*_ |*y*_*t*_ = *i*) be the backward quantity, ∀*i, j*∈ 𝒮_sub_. Initialization (using the sequence-specific None row) is

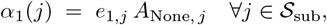

after which the beam *B*_1_ ⊆ 𝒮_sub_ is set to the top-*K* states in *α*_1_, always including None. For *t* ≥ 2,

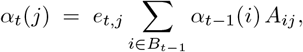

and *B*_*t*_ is updated to the top-*K* states by *α*_*t*_ (again forcing inclusion of None). For the backward pass, set

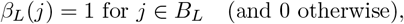

and for *t* ≥ 2,

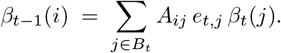

The (beam-restricted) normalizer is

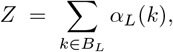

and the posterior marginals are

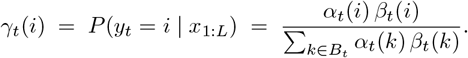

We compute these recursions in log-space for numerical stability. Beam pruning approximates the exact posteriors by restricting all sums to *B*_*t*_. In practice, we use a beam size *K* = 64 and see less than a 0.01% increase in sensitivity on the validation set with a beam size of 128 or higher. See [27] for the classical forward–backward recursions.

#### Maximum expected accuracy (MEA) decoding

Rather than decoding a single most-probable path (Viterbi), which can be brittle when many nearly equivalent paths exist, we use MEA decoding [28]. MEA – also called posterior or optimal-accuracy decoding – uses the posterior marginals from forward–backward and selects the valid path that maximizes expected per-position 0–1 accuracy:

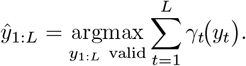

#### Refinement of merged/extended calls

Occasionally, MEA decoding produces calls that are substantially longer than typical for their family, often reflecting merged adjacent domains or boundary over-extension. For each predicted domain, we compute a length ratio (length of the predicted domain divided by the family’s expected length under the learned mid_*f*_ transitions). For calls with length ratio ≥1.5, we perform a refinement step: holding all other decoded calls fixed, we re-decode the corresponding region using a family-restricted 4-state chain None, start_*f*_, mid_*f*_, stop_*f*_ to adjust boundaries and internal structure without permitting family switches. Refinement leaves the non-overlapping domain calls unchanged outside the refined regions.

### Domain Scoring

After MEA decoding and refinement, PSALM outputs domain calls as runs of a family *f* over an interval [*s, u*]. To assign each call a confidence score, we compute (i) a Forward score under a family-restricted 4-state model and (ii) a simple measure of amino-acid frequencies deviation from background, and then combine these with length-based features in a supervised scoring model.

#### Forward score

For a call of family *f* on [*s, u*], we evaluate a 4-state chain {None, start_*f*_, mid_*f*_, stop_*f*_} restricted to this family, using the within-family transitions from *A* and the per-position classifier probabilities restricted to these four states. For the None state we estimate the length-dependent *A*_None,None_ as in the domain-state model, using the called domain’s length ℓ = *u* − *s* + 1. Let *Z*_fam_ be the resulting forward value (sum over all paths) on [*s, u*]. As a null baseline, let *Z*_null_ be the forward value of the same 4-state chain on [*s, u*] when the domain-state probabilities (start/mid/stop) are set to zero at every position while the per-position None probability is retained. We report the Forward score in bits as

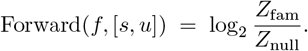

This is analogous to HMMER’s Forward log-odds scoring principle [7] and can also be viewed as a partition-function difference for a fixed linear-chain model [26].

#### Amino-acid composition bias

Let *p*_*k*_ be the empirical frequency of amino acid *k* within the called segment [*s, u*], and let *q*_*k*_ be a fixed background amino-acid frequency computed over all residues in SwissProt release 2025_01 (the background used for scoring; Easel’s `esl-shuffle -G` uses the SwissProt34 background frequencies [29]). The log-likelihood ratio in bits is the KL-divergence of *p*_*k*_ from *q*_*k*_ [30]:

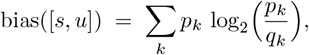

which is zero when the amino acid composition of the called domain matches the background amino acid frequencies.

#### Learning a per-call confidence score

Ideally, we would rank calls using the Forward score alone, since it is the natural segment score induced by our domain-calling model. However, analysis of false positives showed that Forward score alone does not reliably distinguish between true and false positives for short calls (expected or called length ≤50 residues): some short calls (especially from short families or fragmentary/partial hits) score unusually highly relative to true positives of comparable length, and many such false positives also exhibit amino-acid compositions that deviate strongly from background. We therefore train a small supervised scoring model on labeled domain calls that maps per-call features to a confidence score in [0, 1], using the validation set as the source of true-positive labels.

We train a gradient-boosted decision tree model (CatBoost [31], default hyperparameters) as a binary classifier with 3-fold cross-validation over labeled domain calls. Features include: observed length; expected length; absolute length ratio error |1 −len_ratio|; amino-acid composition bias; Forward score; adjusted Forward score (Forward*/* expected length); and domain status (“full” if the decoded path contains both start and stop states for the call, “partial” otherwise). Training labels are constructed from: (i) **false positives**, defined as any domains called on synthetic negative protein sequences generated with Easel using `esl-shuffle -G` (i.i.d. residues drawn from the SwissProt34 background frequencies), with 9M negatives total (1M each at lengths 25, 50, 100, 200, 400, 800, 1600, 3200, and 6400); and (ii) **true positives**, defined as PSALM calls on the validation set (Pfam Full-derived held-out proteins) that double-midpoint overlap a Pfam Full ground-truth domain annotation (the midpoint of the predicted domain lies within the ground truth domain’s boundaries and vice versa). The resulting classifier output is reported as a per-call confidence score and is used to sweep score thresholds for positive domain annotation in benchmarking; filtering by this score is left to the user. We currently do not assign formal statistical significance to individual scores, and instead assess scores empirically in the sensitivity/specificity analyses below.

### Sensitivity and specificity benchmarking

We benchmark PSALM against HMMER for domain detection: recovering true domain intervals with the correct (or closely related) family assignment while controlling false positives. Because Pfam domain calls are defined by HMMER profile HMMs with curated, family-specific gathering thresholds (GAs) [8], benchmarking against GA-filtered Pfam calls alone would largely reflect Pfam’s own HMMER thresholding decisions, making comparison largely circular. We therefore evaluate on a test set in which ground-truth domains are drawn not only from Pfam but also from other InterPro member databases, and are retained only when multiple databases detect the same InterPro domain within broadly agreeing boundaries (similar span/length on the sequence). Regions where member databases disagree substantially on the length or boundary coordinates are treated as ambiguous and excluded from the ground truth (see Benchmark definition).

#### Data overlap and scope of generalization

The two training sets, validation set and the test set are all sourced from UniProt, and may share related sequences, but the training sets and the validation set do not share any identical sequences with the test set. ESM-2 has been pre-trained on a very large sequence database and has already seen many domains and their homologs across our train/test/validation sets. A strict evaluation of remote-homology generalization without information leakage between train and test would require training a protein language model from scratch on carefully held-out sequence sets, which is beyond the scope of this work.

#### Benchmark definition

We derive ground-truth domains for the test set from InterPro [4] in three steps.

a. **Define valid InterPro accessions (IPRs)**. InterPro assigns a shared accession (IPR) to member database models that it considers to represent the same domain family. We keep an IPR only if (i) it includes at least one Pfam family and (ii) its member domain families have similar average lengths: for each member family, we compute its average length over all instances in UniProt, and we require all member families within an IPR to agree within 10% pairwise.
b. **Collect candidate domains per test sequence**. For each test sequence, we keep only the domains with valid IPRs.
c. **Require cross-member database agreement**. If multiple member databases annotate the same IPR on a sequence at overlapping coordinates, we require that all overlapping domains agree by pairwise single-midpoint overlap and do not overlap an already selected domain for the test sequence. If this constraint is met, we keep domain closest to the average length of the IPR. Finally, we construct a non-overlapping set of ground-truth domains by moving from left to right and accepting a domain only if it does not overlap any previously accepted domains.

The resulting test set consists of 88.6M sequences (52B residues; 107.5M domains spanning 14.8B residues). False positives are defined using a separate synthetic negative set of 200,000 sequences whose lengths are sampled from the empirical length distribution of SwissProt release 2021 05, with residues generated i.i.d. from the same SwissProt background amino-acid frequencies. Any predicted domain on these sequences is counted as a false positive. We report specificity as the mean number of false positives per Pfam family, FP*/*family = #{predicted domains on negatives} */*|ℱ|.

#### Match criteria

Predicted domains are compared to ground-truth domains on the same sequence using (i) a positional overlap criterion and (ii) family relatedness. For a ground-truth domain *g* = [*s*_*g*_, *u*_*g*_] and a prediction *p* = [*s*_*p*_, *u*_*p*_], let

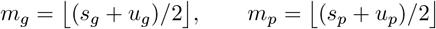

be their midpoints. We define:

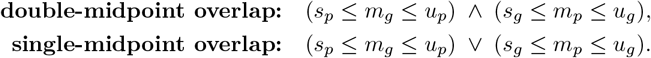

We are primarily investigating whether PSALM can detect signal for a given domain within the ground truth envelope, even if the boundaries of the predicted domain are imprecise. We use single-midpoint overlap as the metric to quantify this detection accuracy. The strict double-midpoint overlap emphasizes a match to the center of the ground truth in addition to signal detection, leading to better quality annotations. We report sensitivity under both midpoint criteria. If a prediction single-overlaps multiple ground-truth domains but does not double-overlap any (e.g., it spans two adjacent ground-truth domains), we credit it as matching exactly one ground-truth domain.

For family identity, a prediction is considered a family match to a ground-truth IPR-labeled domain if its Pfam family is either (i) a Pfam member of that IPR, or (ii) a member of a Pfam clan that appears among the Pfam members of that IPR. Clans are curator-defined groups of homologous Pfam families; all but two IPRs containing multiple Pfam families draw them from a single clan. This allows families within a clan to be credited as detecting the same homologous domain signal. Of the 24k Pfam families, ∼46% are grouped into clans. For all remaining families, an annotation is accurate only when there is an exact match.

#### HMMER comparison

HMMER scores each Pfam family independently and may report overlapping domains on the same sequence, whereas PSALM produces a single, non-overlapping per-position labeling by design. Constructing a non-overlapping set of HMMER hits to mirror PSALM would require discarding some otherwise valid HMMER calls, effectively penalizing HMMER for its ability to report overlapping domains. Instead, for benchmarking we treat a HMMER hit as a true positive whenever it matches a ground-truth domain under the family and overlap criteria above, irrespective of overlaps with other HMMER hits. This favors HMMER, relative to PSALM’s forced non-overlapping labeling, but yields a cleaner comparison of domain detection ability rather than differences in how overlaps are resolved. The comparisons to HMMER are made using profile HMMs that are trained on Training sets 1 and 2 (see Materials and Methods), to match PSALM.

#### Sensitivity–specificity curves

We compare methods by sweeping across score thresholds and plotting the fraction of ground-truth domains recovered (sensitivity) against the mean false positives per Pfam family (Fig. 3). For PSALM, we sweep the learned per-call confidence score from the 0-1 classifier scoring model; for HMMER, we sweep the per-domain E-value. We plot curves under both single- and double-midpoint overlap criteria.

**Figure 3:**
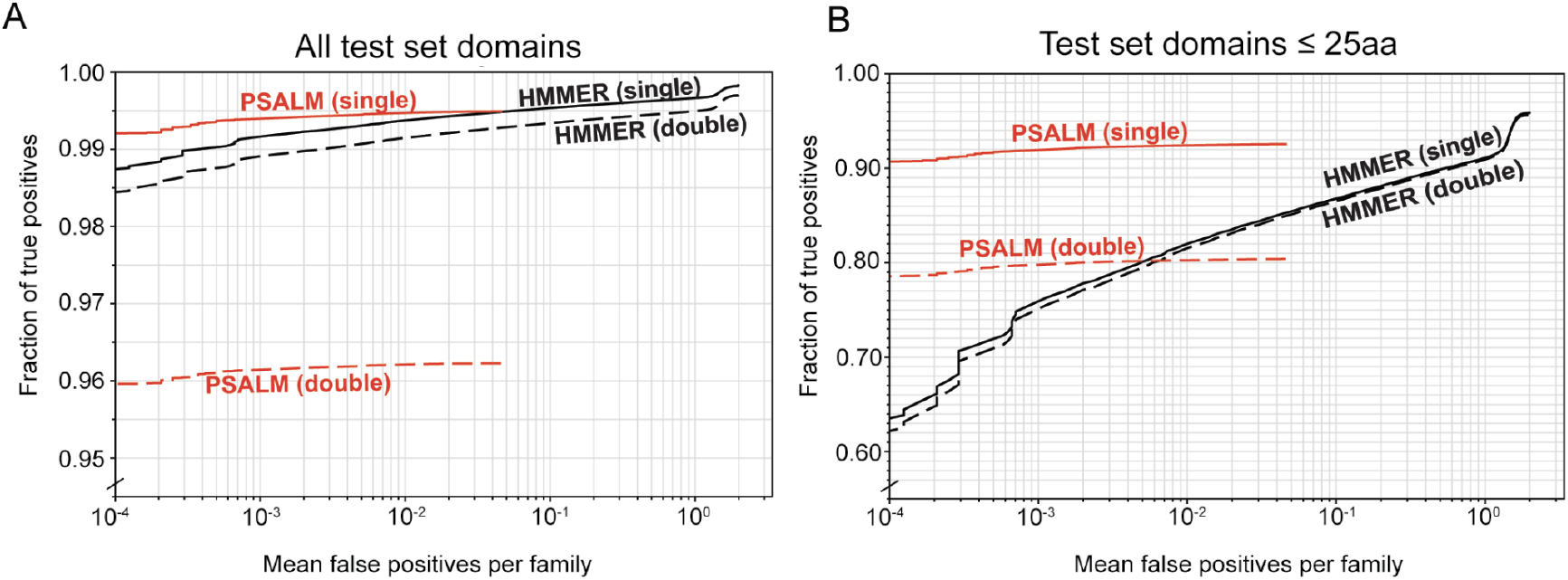
Domain-level sensitivity and false positives per family for PSALM and HMMER. We plot the fraction of true domains recovered versus the mean number of false positives per Pfam family on (**A**) the entire test set and (**B**) only test domains of length at most 25 amino acids. Curves are shown for single- and double-midpoint overlap match criteria.

PSALM achieves HMMER-level sensitivity at a wide range of thresholds, and PSALM’s single-overlap curve exceeds HMMER’s at stringent settings (mean false positives *<* 10^−2^ per family) Fig. 3A. Under the stricter double-midpoint criterion, sensitivity decreases for both methods, reflecting the increased emphasis on matching the centers of test set annotations, with a larger drop in sensitivity for PSALM than for HMMER. For test set domains shorter than 25 amino acids, we show in Fig. 3B that PSALM achieves better sensitivity and specificity than HMMER in both single and double midpoint overlap cases by *∼*25% and *∼*17% respectively at 10^−4^ mean false positives per family.

#### Single vs. double overlap

Discrepancies between single- and double-midpoint overlap fall into two broad categories: over-extension, where a predicted domain call extends beyond the corresponding ground-truth domain, and under-extension, where a predicted domain covers only a portion of the ground-truth domain. PSALM predictions on the test set show that 98% (2.6M domains) of the differences between single and double overlap are over-extensions and the remaining 2% (56K domains) are under-extensions.

To further understand where PSALM achieves good performance using single-midpoint overlap but not double midpoint, we visually inspected examples of over-extensions, two of which are shown in Fig. 4. Figure 4A shows a merged over-extension: where PSALM has incorrectly merged two domains in close proximity into a larger domain — 45% (1.2M domains) of all over-extensions fall into this category. Figure 4B shows an over-extension with respect to the benchmark annotation, which only has one annotated domain. However, the HMMER annotation (profile HMMs built from training sets 1 and 2) identifies not only the benchmark domain but also a second domain, which provides supporting evidence for PSALM’s over-extended domain call. It is likely that this over-extension is another case of a single domain call merging two domains in close proximity.

**Figure 4:**
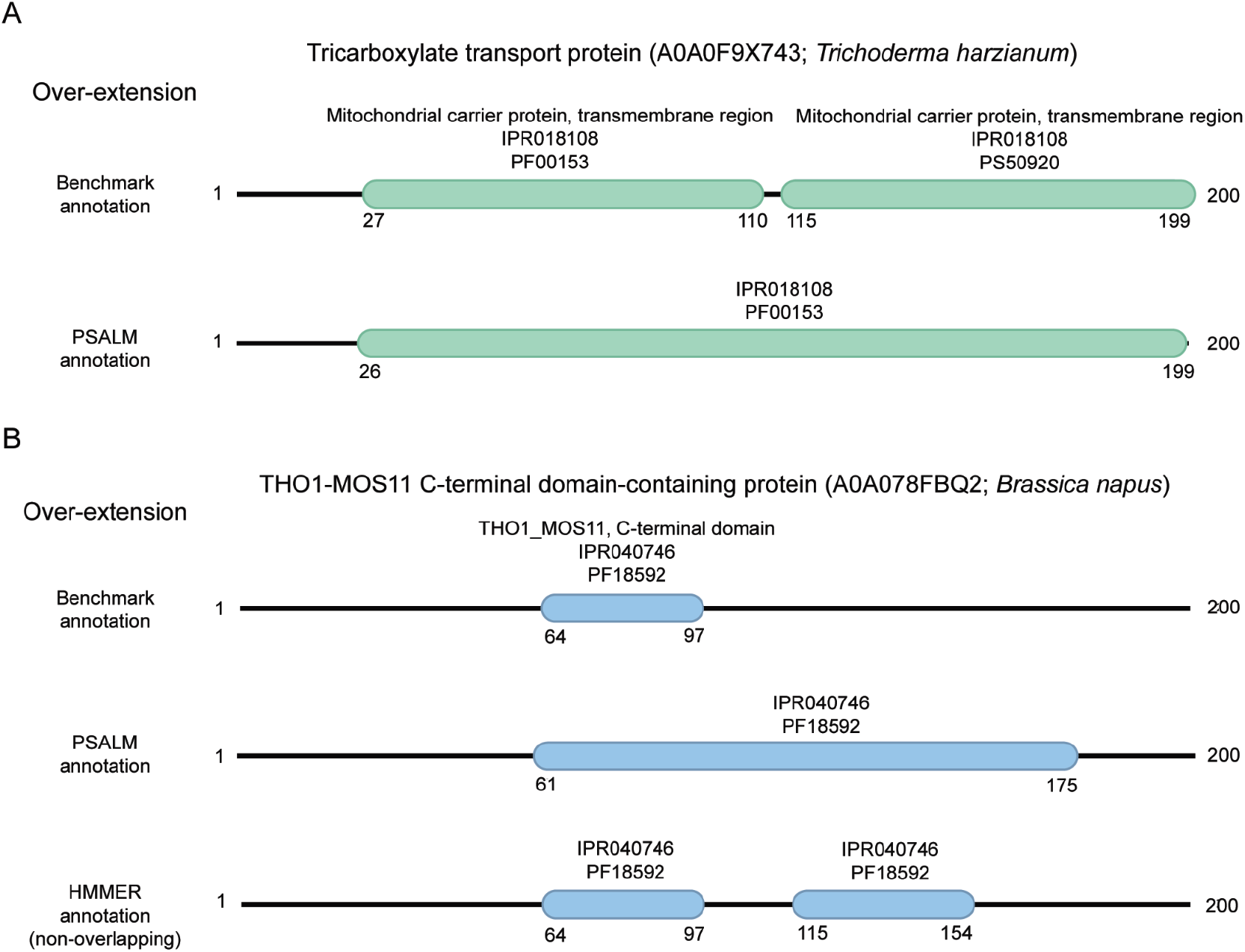
Examples of over-extensions predicted by PSALM. For these two selected sequences, we show both the benchmark annotation and PSALM’s domain calls, as well as the HMMER domain calls for panel B. All annotations and calls are described by their domain family and their sequence start/stop coordinates.

### Coverage on UniProt

Sensitivity and specificity benchmarks quantify agreement with our test set, which is defined primarily by Pfam/HMMER annotations. We also perform a database-wide coverage analysis on UniProtKB 2025 01 [24]. Here, coverage measures (i) the number of sequences for which a method reports at least one domain and (ii) the total number of residues covered by predicted domains.

To make coverage comparable across methods, we report results at matched E-values, the expected number of false-positive hits returned by a search against a database of a given size. Coverage is reported for E-value cutoffs of 0.1, 0.01, and 0.001 expected false positives per Pfam family. On UniProt (253M sequences), these correspond to per-sequence false-positive rates of approximately 4 ×10^−10^, 4 ×10^−11^, and 4 ×10^−12^, respectively. PSALM does not natively compute E-values; it provides a per-domain score from the 0–1 scoring classifier. To place PSALM on the same E-value scale, we estimate an empirical E-value by running PSALM on 30M synthetic negative sequences generated identically to test set negatives and measuring the rate of domains called on these negatives as a function of the score threshold.

HMMER reports higher UniProt coverage than PSALM at the most stringent thresholds (E = 10^−3^ and E = 10^−2^), for both sequence-level and residue-level measures (Table 1). At a more relaxed threshold (E = 0.1), PSALM reports higher sequence and residue coverage than HMMER. Lower E-values correspond to stricter significance thresholds: fewer spurious domain calls are expected, reducing the chance that an incorrect annotation is made and then transferred to newly added sequences to a database due to similarity. The increase in PSALM coverage at E = 0.1 may partly reflect differences in how scores are calibrated. HMMER’s E-values are derived from an explicit statistical model of hit significance and scale with database size, whereas PSALM uses a learned per-domain score that we map to an empirical E-value using synthetic negatives.

**Table 1:** Domain coverage on UniProtKB 2025 01 (253.2M sequences; 91.7B residues) at fixed E-value thresholds.

| E-value | False positive rate | Model | Sequences with $\geq 1$ domain | | Residues in domains | |
| --- | --- | --- | --- | --- | --- | --- |
|  |  |  | # | % of UniProt | # | % of UniProt |
| 0.001 | $3.95 \times 10^{-12}$ | HMMER | 198.6M | 78.4 | 51.6B | 56.2 |
|  |  | PSALM | 156.9M | 62.0 | 45.7B | 49.8 |
| 0.01 | $3.95 \times 10^{-11}$ | HMMER | 201.1M | 79.4 | 52.3B | 57.0 |
|  |  | PSALM | 163.0M | 64.4 | 47.5B | 51.8 |
| 0.1 | $3.95 \times 10^{-10}$ | HMMER | 203.7M | 80.4 | 53.0B | 57.8 |
|  |  | PSALM | 227.6M | 89.9 | 70.6B | 77.0 |

## Discussion

PSALM combines a pretrained protein language model (ESM-2), fine-tuned for per-residue domain-state prediction with a domain-state classifier, with a probabilistic domain decoder to produce non-overlapping domain annotations with explicit boundaries. In domain-level benchmarks, PSALM reaches roughly the same sensitivity–specificity tradeoff as HMMER while making different modeling assumptions: PSALM assigns probabilities over all Pfam families at every residue and then decodes a single non-overlapping annotation for the entire sequence, explicitly weighing competing family hypotheses against each other. This approach may be advantageous in multi-domain proteins and ambiguous regions because selecting one domain family necessarily suppresses alternatives and helps prevent overlapping calls. In contrast, Pfam annotations made using HMMER evaluate one profile HMM per family and reports hits independently across families. These results suggest that a single pLM-based system can be a practical alternative to a large library of per-family models for large-scale protein sequence annotation. Although PSALM relies on a large neural network to produce per-residue domain-state probabilities, final domain calls are obtained by structured inference with an explicit start/mid/stop state model rather than by independent per-position labels alone. For domains shorter than 25 amino acids, which are often repeated in a sequence, the gap between PSALM and HMMER performance suggests that context information from the entire sequence may improve domain detection ability.

Our evaluation has several limitations. First, PSALM does not explicitly model cases where only a portion of the domain family is homologous to the sequence, referred to as fragments. Our state space includes only full start/mid/stop chains, and the transition structure forbids fragments that begin or end in the interior of a domain state chain. The per-residue labels used for training are inferred from Pfam-style annotations that likewise emphasize complete domain instances. As a result, N- and C-terminal fragments or internal domain fragments are either absorbed into the background or counted as over- or under-extended calls relative to the benchmark. Such fragments can still be biologically meaningful, and it may be useful in future work to introduce explicit fragment states or a separate layer of fragment annotation.

We have also not tried to characterize PSALM’s ability to detect remote homologs by creating train and test sets that share little or no similarity with each other. Further, we have not attempted to resolve information leakage from ESM-2 which may be pretrained on sequences from both our train and test sets. This would require a leakage controlled train/test split that is used in the language model pretraining phase as well.

Within this setting, however, PSALM attains high single-midpoint sensitivity at very low false-positive rates on our benchmark. PSALM’s performance shows that residue level embeddings from ESM-2 encode per-position domain family information, which can be recovered by a position-independent neural network (MLP) and smoothed by an HMM-like domain model to predict domains with explicit boundaries. PSALM’s training, inference, and evaluation code are available at https://github.com/Protein-Sequence-Annotation/PSALM, and model weights together with the training/validation/test datasets are available at https://huggingface.co/ProteinSequenceAnnotation.

## Methods

### Data processing and augmentation

#### Truncating long sequences

During training, GPU memory constraints limit the maximum sequence length to *L*_max_ = 4096 amino acids. We therefore slice any training protein longer than *L*_max_ into windows that never split an annotated domain, and re-index domain coordinates relative to each window. This procedure affects only 0.2% of training sequences.

For proteins containing a single annotated domain, we anchor a window to one end of the sequence when possible; otherwise we center it on the domain. For long proteins with multiple domains, we greedily group domains into the minimum number of windows whose span is ≤*L*_max_ (windows may overlap). Each window is suffixed with _`B`/_`M`/_`E` to indicate whether it touches the N-terminus (beginning), neither terminus, or the C-terminus (end) of the parent protein.

For inference and benchmarking in this work, we run PSALM on each input sequence as a whole (no windowing). Users may optionally choose to process long sequences in smaller windows for practical reasons, but this is not required by our evaluation. ESM-2 was pretrained with a maximum context length of 1,024 residues, but it uses rotary positional embeddings (RoPE) [33], so it can be applied to longer sequences at inference time. The accuracy impact of extrapolating beyond the pretraining length is not fully characterized; empirically, in our experiments PSALM produced unchanged domain annotations on sequences up to at least 4,096 residues.

#### Per-residue labels and tokenization

Sequences are tokenized with the ESM-2 tokenizer [15], which converts the amino-acid string into a sequence of discrete vocabulary indices (tokens) expected by the model and inserts special beginning- and end-of-sequence markers (BOS/EOS).

For truncated slices, we keep the usual BOS/EOS convention in the token sequence but control which of BOS and EOS appear using the slice suffix, so that the presence of these markers carries coarse information about where the slice lies within the full-length protein. Slices suffixed with_`B` include BOS but omit EOS;_`E` include EOS but omit BOS; and _`M` omit both, providing coarse information about whether a window comes from the N-terminus, interior, or C-terminus of a longer protein.

### Training

We train the domain-state classifier using a cosine learning-rate schedule with a 2,000-step linear warmup for each augmented training set [34].

#### Model architecture

We use ESM-2 (650M) [15] as the protein language model backbone, implemented with FlashAttention [35, 36]. A three-layer *∼*200M-parameter MLP projects per-residue embeddings (dimension 1,280) to |*S*| = 72,229 domain states with hidden size 2,560:

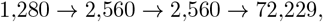

with ReLU activations [37] after the first two linear layers and LayerNorm [38] after each hidden layer. We minimize per-residue cross-entropy between predicted state probabilities and the domain-state labels.

#### Optimization and compute

We use AdamW [39] with (*β*_1_, *β*_2_, *ϵ*) = (0.9, 0.999, 10^−8^), gradient clipping at 1.0, and weight decay 0.01 on ESM-2 parameters (0 for the MLP head). Training uses an effective batch size of 65,568 tokens (up to 16 sequences of length 4096+2 tokens per step, including BOS/EOS where present). All experiments were run on a single 4 *×*NVIDIA H100 80GB node; training completes in ∼2.5 days.

### Inference

#### Cross-family transitions

To allow plausible transitions between different families that may not appear in the supervised training labels, we add a small amount of probability mass to filtered classes of cross-family transitions and redistribute it over transitions that are unobserved in the training corpus. For each source row considered, we distribute the corresponding mass uniformly across cross-family destinations that are unobserved in the training corpus (i.e. transitions with *A*_*ij*_ = 0 before adding these terms), then renormalize the row:

- **Mid** → **(unobserved) cross-family starts**. For each middle state *i* = mid_*f*_ and each start state *j* = start_*g*_ with *g* ≠ *f* such that the transition is unobserved in the training corpus (*A*_*ij*_ = 0), add

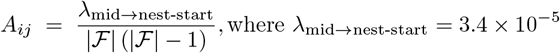

then renormalize row *i*.
- **Stop** → **(unobserved) cross-family middles**. For each stop state *i* = stop_*f*_ and each middle state *j* = mid_*g*_ with *g*≠ *f* such that the transition is unobserved (*A*_*ij*_ = 0), add

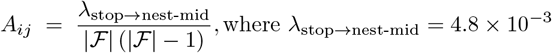

then renormalize row *i*.
- **Stop** → **(unobserved) starts**. For each stop state *i* = stop_*f*_ and each start state *j* = start_*g*_ (including *g* = *f*) such that the transition is unobserved (*A*_*ij*_ = 0), add

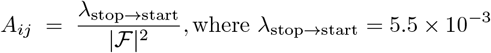

then renormalize row *i*.

All edges already present in *A* are retained. Each updated row *i* is normalized so that S_*j*_ *A*_*ij*_ = 1.

#### Refinement of merged/extended calls

For each decoded call of family *f* spanning [*s, u*], we compute a length ratio

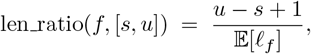

where 𝔼 [ℓ_*f*_] is the expected length of a single *f*-domain implied by the within-family mid_*f*_ exit probability. Using the within-family transitions, 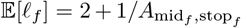 (start + expected mid-run + stop). We refine only calls with len_ratio ≥1.5.

Refinement is performed independently per selected call. All decoded calls not under refinement are treated as preserved: during refinement of [*s, u*], we prevent overlap with preserved calls by masking perposition classifier scores so that preserved positions assign all probability mass to None (equivalently, all non-None states receive score 0). Positions outside preserved spans retain their original scores.

We then re-decode using a family-restricted 4-state chain {None, start_*f*_, mid_*f*_, stop_*f*_} that forbids family switches. The transition structure is the within-family submodel induced by *A*: domains may start only via None → start_*f*_, start_*f*_ transitions only to mid_*f*_, mid_*f*_ either self-loops or exits to stop_*f*_, and stop_*f*_ returns to None or start_*f*_ (with the corresponding probabilities from *A*, restricted and renormalized to these four states). We run forward–backward on this 4-state model and decode with MEA. The resulting refined path replaces the original call of family *f* in that region; preserved calls and all regions outside refined spans are unchanged.

## Acknowledgments

We thank Tom Jones for discussions on nested domains and advice on identifying challenging domain annotation examples. High-performance computational resources were provided by Harvard FAS Research Computing.

Research reported in this publication was supported in part by the National Human Genome Research Institute of the US National Institutes of Health. The content is solely the responsibility of the authors and does not necessarily represent the official views of the National Institutes of Health.

## Author Contributions

Conceived and designed the experiments: AS, KK, SRE. Performed the experiments/developed methods: AS, KK. Analyzed the data: AS, KK. Wrote the paper: AS, KK, SRE.

## Notes

https://github.com/Protein-Sequence-Annotation/PSALM

https://huggingface.co/collections/ProteinSequenceAnnotation/psalm-2

https://pypi.org/project/protein-sequence-annotation/

### Competing Interest Statement

The authors have declared no competing interest.

### Summary of Updates

This version of the manuscript has been revised to update data curation, methodology, and results

